# Single-nucleotide and Copy-number variance related to severity of Hypospadias

**DOI:** 10.1101/282459

**Authors:** Neetu Singh, Devendra Kumar Gupta, Shilpa Sharma, Dinesh Kumar Sahu, Archana Mishra, Devendra Yadav, Jiledar Rawat, Arun K Singh

**Affiliations:** Molecular Biology Unit (Center for Advanced Research), King George’s Medical University, Lucknow, India-226 003; Department of Pediatric Surgery, All India Institute of Medical Sciences, New Delhi, India-110029; Department of Plastic Surgery, King George’s Medical University, Lucknow, India-226 003

**Keywords:** Hypospadias, Copy Number Variations, PRSS3P2, TSC1, DMBT1, STS, SRD5A2, STARD3

## Abstract

The genetic association of Hypospadias-risk studies has been conducted in Caucasians, Chinese-Han populations and few in Indian populations. Although no comprehensive approach has been followed to assess genetic involvement in the severity of the disorder. The study evaluated to establish the correlation between genotyped SNPs/CNVs and Hypospadias-severity by an association in a total 30 SNPs in genes related to sex hormone-biosynthesis and metabolism; embryonic-development and Phospholipase-D-signalling pathways on 138 surgery-confirmed hypospadias-cases from North-India (84 Penile and 28 cases of Penoscrotal-Hypospadias compared against 31 cases of Glanular+Coronal), and analyzed and identified copy number variants (CNVs) in four Familial samples (18 members) and three paired-sporadic cases (6 samples) using array-based comparative-genomic-hybridization and validated in 32 Hypospadias samples by TaqMan assay. Based on Odds Ratio at 95% CI, Z Statistic and Significance Levels, STS gene-rs17268974 was associated with Penile-Hypospadias and 9-SNPs (seven-SNPs (rs5934740; rs5934842; rs5934913; rs6639811; rs3923341; rs17268974; rs5934937) of STS gene; rs7562326-SRD5A2 and rs1877031-STARD3 were associated with Penoscrotal-Hypospadias. On aggregate analysis with p <0.001, we identified homozygous-loss of Ch7:q34 (PRSS3P2, PRSS2). On validation in previously CNV-characterized and new (32-hypospadias-cases), we identified PRSS3P2-loss in most of the grade 3 and 4 hypospadias. Hence, Grade 1 and 2 (coronal and granular) show no-PRSS3P2-loss and no-association with SNPs in STS; SRD5A2; STARD3-gene but Grade 3 and 4 (Penile and Penoscrotal) show PRSS3P2-loss accompanied with the association of SNPs in STS; SRD5A2; STARD3. Hence, homozygous-loss of PRSS3P2 accompanied with the association of STS; SRD5A2; STARD3 may link to the severity of the disease.

## Introduction

Hypospadias is a common congenital malformation in which the urethral meatal opens on the ventral surface of the penis and is often associated with cryptorchidism and disorders of sexual differentiation (Klip et al. 2002). Prevalence of Hypospadias varies considerably across countries ranging from 1 in 125 to 250 male live births. Embryological studies have demonstrated that, depending on the stage at which the urethral development arrests, the meatal opening can be anywhere along the shaft of the penis or, in more severe forms, within the scrotum or in the perineum. Hence, the question arises what is the underlying cause of the severity of common anomaly of the male genitalia? This may be due to the genetic variations, however, the genomic regions responsible for hypospadiac pathogenesis remains poorly understood. Also, there are some reports in decipher database which show that these aberrations may be present in the autosomes or sex-linked chromosomes (Harrison et al. 2013; de la Chapelle 1987). Till date, both population based case-control and genome-wide association studies have shown some genes to be associated with hypospadias. Most assessed Population-based: Case-Control Association Studies like Diacylglycerol Kinase Kappa (*DGKK*6)-converts diacyl glycerol to phosphatidic acid-rs4554617, Master Mind Like domain (MAMLD1) mutations/inframe splice variants generating dysfunctional proteins and/or unstable mRNAs (Liu et al. 2017); Steroid 5-alpha reductase (SRD5A2-V89L) - Conversion of testosterone to dihydrotestosterone (Samtani et al. 2011), Steroid 5-alpha reductase (SRD5A2-A49T, SRD5A2-R227Q) and TA repeat gene polymorphisms (Samtani et al. 2015), MAMLD1-activates promoter of non-canonical NOTCH target Hes 3 promoter (Novel-P299A misense and c.2960C>T in in 3’ UTR of Exon 7; (Ratan et al. 2016), CYP17A1-Conversion of pregnenolone and progesterone (Samtani et al. 2010; Qin et al. 2012), Hydroxy Steroid 17-b dehydrogenase (HSD3B2)-Interconversion of androgens and progesterone related hormones (Codner et al. 2004), Hydroxy Steroid 17-b dehydrogenase HSD17B3-Interconversion of oestrogens and androgens (Sata et al. 2010), CYP1A1-Contributes to 2-alpha hydroxylation of oestrogens (Kurahashi et al. 2005), Androgen receptor gene CAG repeat polymorphism and risk of isolated hypospadias (Huang et al. 2015) and Tissue-specific roles of FGF signaling in external genitalia development. (Harada et al. 2015).

Independent association of 18 genomic regions has been shown to be associated with Hypospadias through the most important study of Denmark. This was performed on 1,006 surgery-confirmed hypospadias cases and 5,486 controls from Denmark and validation in 1,972 cases and 1,812 controls from Denmark, the Netherlands and Sweden showed an independent association of 18 genomic regions with *P* < 5 × 10^−8^ with the development of Hypospadias. The identified regions included genes with key roles in embryonic development (including *HOXA cluster (HOXA4 and HOX A13), IRX5*, *IRX6,* and *EYA1*) (Geller et al. 2014). Carmichael *et al.,* (Carmichael et al. 2014) also examined the association of variants of genes related to sex hormone biosynthesis and metabolism in the different type of 633 cases of Hypospadias (84 mild, 322 moderates, 212 severe and 15 undetermined severity) and 855 population-based non-malformed male controls born in California from 1990 to 2003. They examined 332 relatively common tag single-nucleotide polymorphisms (tagSNPs) in 20 genes. Of these 332, several significant SNPs were identified one in CYP3A4, four in HSD17B3, one in HSD3B1, two in STARD3, 10 in SRD5A2 and seven in STS which showed association with hypospadias risk. All these genes contributed to sex hormone biosynthesis and metabolism.

Based on above International and National studies, a total of 30 SNPs in different genes related to sex hormone biosynthesis and metabolism [CYP3A4 one SNP-rs12333983; HSD17B3 four SNPs-rs12552648; rs8190566; rs8190557; rs2026001; STARD3 two SNPs-rs1874224; rs1877031; SRD5A2-ten SNPs-rs1042578; rs9332975; rs2281546; rs28383032; rs6543634; rs2268794; rs725631; rs7562326; rs765138; rs519704; STS seven SNPs-rs5934740; rs5934842; rs5934913; rs6639811; rs3923341; rs17268974; rs5934937)]; embryonic development (*HOXA4*-rs1801085, *IRX5*-rs17208368, *IRX6-*rs6499755, *ZFHX3* (intronic)-rs1858800 and *EYA1* rs16937456) and Phospholipase D signaling pathways [rs4554617 of DGKK (Diacylglycerol Kinase Kappa) which phosphorylates diacylglycerol (DAG) to generate phosphatidic acid (PA)] have been identified which needs to be evaluated with the severity of disease.

We hypothesized to identify variants in genes related to sex hormone biosynthesis and metabolism; embryonic development and Phospholipase D signaling pathways on blood of 138 surgery-confirmed hypospadias cases from North India to establish a correlation between genotyped SNPs and severity of Hypospadias. We also analyzed and identified copy number variants (CNVs) in Four Family samples-18 members and 3 paired-sporadic cases (6-samples) using array-based comparative genomic hybridization-Cytoscan 750K array and validated identified CNVs in 32 Hypospadias samples-TaqMan assay using the probe against target sequence: PRSS3P2 and reference sequence TERT.

## Methods

### Ethical approval

This study was approved by the Institutional Review Board of the King George Medical University (Lucknow, India) and patient consent was obtained for blood collection as well as for the release of all medical records.

### Patient Selection for Association Study

The study population includes 138**-**cases classified on the basis of severity of hypospadias, based on the reported anatomical position of the urethral opening before surgery. Cases for which the anatomical position (Mild cases-those for which the meatus will be limited to the coronal or granular penis; Moderate cases-those for which the meatus will be on the penile shaft; Severe cases-those for which the meatus will be at the penoscrotal junction, scrotal or perineal area) were included in the study. Cases for which the anatomical position was not described and Cases having a known single gene disorder or chromosomal abnormality were excluded from the study.

The studied genetic variance related to cause and severity of Hypospadias were as following (a) sex hormone biosynthesis and metabolism: CYP3A4 one SNP-rs12333983; HSD17B3 four SNPs-rs12552648; rs8190566; rs8190557; rs2026001; STARD3 two SNPs-rs1874224; rs1877031; SRD5A2-ten SNPs-rs1042578; rs9332975; rs2281546; rs28383032; rs6543634; rs2268794; rs725631; rs7562326; rs765138; rs519704; STS seven SNPs-rs5934740; rs5934842; rs5934913; rs6639811; rs3923341; rs17268974; rs5934937); (b) embryonic development pathway including homeobox gene family (*HOXA4*-rs1801085, *IRX5*-rs17208368, *IRX6*-rs6499755, *ZFHX3* (intronic)-rs1858800 and *EYA1-*rs16937456); and (c) Phospholipase D signaling pathway [rs4554617 of DGKK (Diacylglycerol Kinase Kappa) which phosphorylates diacylglycerol (DAG) to generate phosphatidic acid (PA)].

### Patient selection-family-based study for Copy Number Variations

We also assessed four Family samples (18 members) and three paired-sporadic cases (6-samples) using array-based comparative genomic hybridization-Cytoscan 750K array (Table 1) and validated identified CNVs in 32 Hypospadias samples-TaqMan SNPs. The patients recruited for Hypospadias surgery in the Department of Plastic and Pediatric Surgery at King George Medical University in the year 2013-14 were taken for this study. Importantly, the collected samples were from unrelated families. As designed above, in discovery phase four families (18 samples) and in validation phase three affected individuals and their fathers (6 samples) were considered as shown in Table 1., 20 new samples were collected for validation, from All India Institute of Medical Sciences, New Delhi (AH01-AH-12) and King George Medical University (Lucknow, India) (H11, H13, H15, H16-18, and H33-34). More, CNV characterized 12-samples (H1, H2, H7, H9, H19, H20, H21, H22, H27, H32, H37, and H38) were also considered for validation.

**Table 1:** Details of four families considered for discovery phase and three paired-sporadic cases of validation phase.

| Discovery phase | Family 1 | Family 2 | Family 3 | Family 4 |
| --- | --- | --- | --- | --- |
|  | Affected twins-H1-(Sub-coronal)<br>Affected twins-H2-(Glanular)<br>Mother-H3<br>Father- H4 | Affected Son-H19-(Sub-coronal)<br>Affected Son-H20-(Proximal Penile)<br>Affected Son-H21-(Proximal Penile)<br>Affected Son-H22-(Proximal Penile)<br>Non-Affected Son- H23<br>Mother-H24<br>Father- H25 | Affected Son-H27-(Sub-coronal)<br>Mother-H28<br>Father- H29 | Affected Son-H37-(Proximal Penile)<br>Affected Son-H38-(Penoscrotal)<br>Mother-H39<br>Father- H40 |
| Sporadic phase | Sporadic Case 1 | Sporadic Case 2 | Sporadic Case 3 |  |
|  | Affected Son-H7-(Penoscrotal-with Chordee)<br>Father- H8 | Affected Son-H9-(Proximal Penile)<br>Father- H10 | Affected Son-H32-(Proximal Penile)<br>Father- H35 |  |

### Isolation of Genomic DNA

Fresh, peripheral blood samples were collected from 138 surgery confirmed cases for association study and from hypospadias patient and its family for CNV study were processed for DNA isolation and quality check. Briefly, Genomic DNA was isolated from blood samples using QIAGEN DNA purification kit (QIAGEN, Hilden, Germany) according to the manufacturer’s instructions. Isolated DNA was checked for quality on an agarose gel, quantitated by the spectrophotometer, and finally stored at −20°C for further use.

### SNP analysis through MassArray Analyzer

For 138 surgery-confirmed hypospadias cases, locus-specific primers were designed by Assay Design Suite (ADS) and region of interest was PCR amplified. The iPLEX extension primers for each SNP locus were annealed to amplified DNA fragments and extended by iPLEX single base extension mechanism to identify the genotyping target site. The SNP alleles (homozygous or heterozygous) were analyzed by MassArray Analyzer via an ApectroCHIP Array (a silicon chip with pre-dispensed matrix crystal) and “Typer software”, an integrated data analysis tool.

### CytoScan 750K Array for copy number analysis and Gene Identification

The Genome-Wide Human CytoScan 750K Array (Affymetrix, CA, USA) was used to analyze genomic alterations according to the manufacturer’s protocol. Briefly, 250 ng of genomic DNA was isolated from peripheral blood of an affected individual and his parents. Isolated DNA was fragmented through the restriction enzyme NspI and then adapter was ligated to fragmented DNA. The process of ligation was followed by PCR amplification using the NspI-adapter-specific primer. Further, the quality of the amplified product was checked on a 2% TBE gel. After confirming that the products were between 150 and 2000 bp in length, the amplified products were subjected to purification through magnetic beads (Affymetrix). The quantity of the PCR product were measured by Quawell spectrophotometer, the product more than 3μg were considered for fragmentation using DNase I and visualized on a 4% TBE agarose gel to confirm that the fragment size ranged from 25 to 125 bp. Subsequently, the fragmented PCR products were end-labeled with biotin and hybridized to the array for 17 hours. Arrays were then washed and stained using a GeneChip® Fluidics Station 450 and scanned using an Affymetrix GeneChip® Scanner 3000 7G. Scanned data files were generated using Affymetrix GeneChip Command Console Software, version 1.2, and analyzed with Affymetrix® Chromosome Analysis Suite v1.2 (ChAS) (Affymetrix Inc., USA) and Nexus copy number 7.0 software. Thresholds of log2 ratio with default parameters were used to categorize altered regions as CNV gains (amplification) and copy number losses (deletions), respectively. Amplifications and deletions were analyzed separately. To exclude aberrations representing common normal CNVs and identify disease-related CNVs, all the identified CNVs were compared with those reported in the Database of Genomic Variants (DGV, http://projects.tcag.ca/variation/) and decipher databases respectively.

### Validation of the copy number variation by TaqMan copy number assay

TaqMan® Copy Number Assays is a process to detect and measure copy number variation within the human genomes using TERT as a reference in a duplex real-time polymerase chain reaction (PCR). The primers and probes are synthesized both against the target gene/genomic sequence (FAM™ dye-labeled MGB probe) and the reference sequence (VIC® dye-labeled TAMRA™ probe). This method measures the CT difference between target and reference sequences and compares CT values of test samples to a calibrator sample(s) known to have two copies of the target sequence. Briefly, 20ng of purified genomic DNA is processed both with the TaqMan® copy number assay kit, containing two primers and a FAM™ dye-labeled MGB probe to detect the PRSS3P2 and the TaqMan® copy number reference assay, containing two primers and a VIC® dye-labeled TAMRA™ probe to detect the genomic DNA reference sequence-TERT. After adding the TaqMan® Genotyping Master Mix, (AmpliTaq Gold® DNA Polymerase, and dNTPs) real-time PCR reaction is carried out under universal thermal cycling condition i.e. denaturation at 95 °C for 10 min followed by 40 cycles at 95 °C 15 sec and 60 °C for 60 sec. The data obtained is analyzed on CopyCaller™ Software v1.0 for measuring the copy numbers of the PRSS3P2 genes in 32 hypospadias samples.

### Statistical analysis

Data were statistically analyzed by using appropriate statistical package for the study (SPSS). Groups of continuous variables will be compared by student’s t-test or analysis of variance (ANOVA), while discrete (categorical) data will be analyzed with the chi-square test. Odds ratio will be calculated between cases and controls.

## Results

Samples with approx. 100% call rate were genotyped through “Typer software”. Both, 84 Penile (14-Proximal Penile; 21-Mid Penile and 48-Distal Penile) and 28 cases of Penoscrotal Hypospadias were compared with 31 cases of Glanular+Coronal for all 30 SNPs through Odds Ratio at 95% CI, Z Statistic, and Significance Levels. One SNP-Steroid Sulfatase-STS gene-rs17268974 (Microsomal) was associated with Penile type of Hypospadias (Table 2a), while nine SNPs including all the seven SNPs (rs5934740; rs5934842; rs5934913; rs6639811; rs3923341; rs17268974; rs5934937) of STS gene; rs7562326 of SRD5A2 (Steroid 5-alpha reductase) and rs1877031 of STARD3 gene-STAR-related lipid transfer were associated with Penoscrotal type of Hypospadias (Supplementary Table 1 and Table 2b).

**Table 2: (a).** Significant association as evidenced by Odds Ratio at 95% CI, Z Statistic and Significance Levels of rs17268974-Steroid Sulfatase-STS gene with Penile type of Hypospadias. For genotyping we compared 31 Glanular+Coronal vs. 84 Penile (proximal, Mid and distal) cases through “Typer software”-Sequenom.

| Number of Samples | Type of Hypospadias (Comparison between Penile and Glanular plus coronal) | rs17268974 STS |
| --- | --- | --- |
| 14 | PROXIMAL PENILE |  |
|  |  | TT-11 |
|  |  | AA-3 |
| 21 | MID PENILE |  |
|  |  | TT-18 |
|  |  | AA-3 |
| 48 | DISTAL PENILE | TA-1 |
|  |  | TT-37 |
|  |  | AA-10 |
| 84 | TOTAL PENILE | TA-19 |
|  |  | TT-66 |
|  |  | AA-16 |
|  | Penile Region (84) | AA+TA=35 |
|  |  | TT-66 |
|  | Glanular and Coronal (31) | AA=3 |
|  |  | TT-28 |
|  | ODDS RATIO | 4.9495 |
|  | 95%CI | 1.4050 to 17.4359 |
|  | Z STATISTIC | 2.489 |
|  | SIGNIFICANCE LEVEL | P = 0.0128 |

**Table 2: (b).** On comparing 31 Glanular+Coronal vs. 28Penoscrotal, significant association of nine SNPs including all the seven SNPs (rs5934740; rs5934842; rs5934913; rs6639811; rs3923341; rs17268974; rs5934937) of STS gene; rs7562326 of SRD5A2 (Steroid 5-alpha reductase) and rs1877031 of STARD3 gene with Penoscrotal type of Hypospadias was identified.

| Number of Samples | Type of Hypospadias (Comparison between Penoscrotal and Glanular plus coronal) | Rs17268974 (STS) | Rs1877031 (STARD3) | Rs3923341 (STS) | Rs5934740 (STS) | Rs5934842 (STS) | Rs5934913 (STS) | Rs5934937 (STS) | Rs6639811 (STS) | Rs7562326<sup>6</sup> (SRD5A2) |
| --- | --- | --- | --- | --- | --- | --- | --- | --- | --- | --- |
| 28 | PENOSCROTAL | TA-4 | AG-16 | TC-4 | AG-4 | CA-4 | AG-4 | GC-4 | GA-4 | CT-7 |
|  |  | TT-15 | AA-7 | CC-15 | GG-15 | CC-15 | GG-15 | CC-15 | AA-15 | TT-21 |
|  |  | AA-9 | GG-5 | TT-9 | AA-9 | AA-9 | AA-9 | GG-9 | GG-9 |  |
|  |  | TA+AA=13 | AG+GG=21 | TC+TT=13 | AG+AA=13 | CA+AA=13 | AG+AA=13 | GC+GG=13 | GA+GG=13 |  |
| 31 | GLANULAR+CORONAL |  | AG-9 |  |  | CA-1 |  |  | GA-1 |  |
|  |  | TT-28 | AA-18 | CC-28 | GG-28 | CC-27 | GG-28 | CC-28 | AA-27 | TT-31 |
|  |  | AA-3 | GG-4 | TT-3 | AA-3 | AA-3 | AA-3 | GG-3 | GG-3 |  |
|  |  | AA=3 | AG+GG=13 | TC+TT=3 | AG+AA=3 | CA+AA=4 | AG+AA=3 | GC+GG=3 | GA+GG=4 | CT+CC=0 |
|  | ODDS RATIO | 8.0889 | 4.1538 | 8.0889 | 8.0889 | 5.85 | 8.0889 | 8.0889 | 5.85 | 21.9767 |
|  | 95%CI | 1.9881 to 32.9116 | 1.3637 to 12.6529 | 1.9881 to 32.9116 | 1.9881 to 32.9116 | 1.6165 to 21.1711 | 1.9881 to 32.9116 | 1.9881 to 32.9116 | 1.6165 to 21.1711 | 1.1915 to 405.3630 |
|  | Z STATISTIC | 2.92 | 2.506 | 2.92 | 2.92 | 2.692 | 2.92 | 2.92 | 2.692 | 2.078 |
|  | SIGNIFICANCE LEVEL | P = 0.0035 | P = 0.0122 | P = 0.0035 | P = 0.0035 | P = 0.0071 | P = 0.0035 | P = 0.0035 | P = 0.0071 | P = 0.0377 |

Through Nexus Copy number, we performed different levels of computation based on the relative probe intensities between affected and normal individual. Firstly, we analyzed the individual family and paired-sporadic cases (hypospadias patient and his father) (Table 1), to identify the common regions of gain or loss (segmented region of gain or loss) in pro-band compared to parents in familial cases and father in sporadic cases. In the next level, we combined all the results into three groups; I group including all hypospadias samples, II group including all mother samples and III group including all father samples and identified the minimal region of overlap among cases and control (parental) samples. The identified regions of common aberrations gave us the lead to identify the genes and biological pathway being interrupted as well as also gave the information regarding the inheritance pattern of common aberrations.

On analyzing individual family/paired-sporadic cases (Table 1), we observed sharing of Chr6:p25.3-gain (DUSP22), among 75% members of the family I. In second family Ch5:q11.1-gain (PARP8), Ch20:p11.1-gain (MIR663AHG, MIR663A), Ch7:q34-loss (PRSS3P2, PRSS2), Ch10:q26.13-loss (DMBT1) occurred with the frequency of 42.86, 42.86, 57.14 and 42.86 respectively (Supplementary table2a-c). In the third family, no significant CNVs were observed (Supplementary table3a-d). In fourth family gain of Ch22: q11.23 - q12.1 (IGLL3P, LRP5L, CRYBB2P1, MIR6817), gain of ChY: q11.222 (FAM224A, FAM224B, FAM41AY1, FAM41AY2, HSFY2, HSFY1, TTTY9A, TTTY9B, TTTY9A, TTTY9B, HSFY1, HSFY2, TTTY14, CD24) and loss of Ch3:p11.1 - q11.1 (no gene) was observed with the frequency of 75% among the family members (Supplementary table4a-e). Next, affected sporadic sample H7 and related father H8 showed no significant CNVs (Supplementary table5a-c). However, affected sample H9 and related father H10 reported Ch3:p11.1 - q11.1-loss (no gene) and Ch7:q34-loss (PRSS3P2, PRSS2) with a frequency of 50 and 100% respectively (Supplementary table2 6a-c). Additionally, in affected sample H32 and related father H35, Ch4:q35.1 - q35.2-gain (TLR3, FAM149A, FLJ38576, CYP4V2) and Ch7:q34-loss (PRSS3P2, PRSS2) was observed with a frequency of 100 and 66.66% respectively (Table 2a and Supplementary Table 7).

When analyzed all the Hypospadias samples we identified loss of Ch3:p11.1 - q11.1 (no gene), Ch7:q34 (PRSS3P2, PRSS2) and Ch10:q26.13 (DMBT1) with the frequency of 36.36%, 54.55%, and 36.36% respectively. Additionally, gain of Ch9:q34.13 (TSC1) and Ch10:q11.21 (RET) was observed with the frequency of 36.36%. When compared to all mother samples we identified loss of Ch3:p11.1 - q11.1 region (no gene-50% frequency) was common to Hypospadias cases. While in comparison with all father samples, Ch7:q34-loss (PRSS3P2, PRSS2) and Ch10:q26.13-loss (DMBT1), was common to Hypospadias samples. However, gain of Ch9:q34.13 (TSC1) and Ch10:q11.21 (RET) was de-novo to Hypospadias samples (Table 3a).

**Table 3a:** Analysis of family, paired-sporadic and Mother, Father and Hypospadias cases through Nexus copy number software version 7. P value cut off 0.05 and aggregate percent cut off was 35%

| Samples | Region | Region Length | Cytoband Location | Event | Genes | Frequency % | P-Value | % of CNV Overlap | B/P Genes | Gene Symbols | segmental_dups.bed |
| --- | --- | --- | --- | --- | --- | --- | --- | --- | --- | --- | --- |
| I Family | chr6:252,123-331,282 | 79159 | p25.3 | CN Gain | 1 | 75 | 0 | 100 |  | DUSP22 |  |
| II Family | chr5:50,016,088-50,045,767 | 29679 | q11.1 | CN Gain | 1 | 42.85714286 | 0 | 100 | PARP8 | PARP8 |  |
|  | chr20:26,129,788-26,266,313 | 136525 | p11.1 | CN Gain | 2 | 42.85714286 | 0 | 100 |  | MIR663AHG , MIR663A | chr7:57582063, chr4:33848843, chr4:33823557, chr4:34428591, chr9:69891706, chr10:27604109, chr10:45680038, chr10:37424225, chr18:14759247, chr10:45651782, chr3:76014707, chrUn_gl000220:105861, chrUn_gl000220:149833, chr3:75976115, chr3:75920589, chr22:17225458, chr7:57714356, chr17:21895809, chr9:44817871, chr9:45049257, chr9:45784012, chr9:46170362, chr9:46330170, chr20:29572484, chr20:25850889, chr7:57873079, chr20:25781227, chr20:26031728, chr20:29419569, chr15:47398229, chr19:37524431, chr12:37987839, chrUn_gl000246:31380, chr7:57719944, chr9:67729475, chr9:44808260, chr9:45052984, chr9:45787738, chr9:46174087, chr9:46320568, chr19:37526420, chr17:21883168, chr22:17234097, chr20:26018224, chr7:57861338, chr20:29575544, chr19:40152920 |
|  | chr7:142,476,116-142,486,098 | 9982 | q34 | CN Loss | 2 | 57.14285714 | 0 | 100 |  | PRSS3P2, PRSS2 | chr7:142457676, chr9:33611677 |
|  | chr10:124,338,004-124,382,194 | 44190 | q26.13 | CN Loss | 1 | 42.85714286 | 0 | 100 | DMBT1 | DMBT1 |  |
| III Family | - | - | - | - | - | - | - | - | - | - | - |
| IVFamily | chr22:25,691,453-25,909,031 | 217578 | q11.23 - q12.1 | CN Gain | 4 | 75 | 0 | 100 |  | IGLL3P, LRP5L, CRYBB2P1, MIR6817 | chr22:25976766, chr22:26000004, chr22:23914480, chr22:23973678, chr22:23964180, chr22:23942032, chr11:68170544, chr11:68124143, chr11:68071002, chr22:25672707, chr22:25955540, chr22:25971578, chr22:23936734, chr22:25622856, chr22:25649610 |
|  | chrY:20,455,205-21,203,264 | 748059 | q11.222 | CN Gain | 14 | 75 | 0 | 97.791217 | CD24, TTTY14 | FAM224A, FAM224B, FAM41AY1, FAM41AY2, HSFY2, HSFY1, TTTY9A, TTTY9B, TTTY9A, TTTY9B, HSFY1, HSFY2, TTTY14, CD24 | chrY:19567683, chr12:38469473, chr16:90151279, chr3:197840878, chr12:34293493, chrUn_gl000234:3688, chrUn_gl000232:14927, chr12:34274694, chrY:20841912, chrY:24194698, chrY:24682840, chrY:24194698, chrY:24682840, chrY:20612098, chr15:56363536 |
|  | chr3:90,310,894-91,000,000 | 689106 | p11.1 - q11.1 | CN Loss | 0 | 75 | 0 | 28.14661315 |  |  | chr3:93587959 |
| H7 and H8 | - | - | - | - | - | - | - | - | - | - | - |
| H9 and H10 | chr3:90,310,894-91,000,000 | 689106 | p11.1 - q11.1 | CN Loss | 0 | 50 | 0 | 28.14661315 |  |  | chr3:93587959 |
|  | chr7:142,476,116-142,486,098 | 9982 | q34 | CN Loss | 2 | 100 | 0 | 100 |  | PRSS3P2, PRSS2 | chr7:142457676, chr9:33611677 |
| H32, 35 and 36 | chr4:186,936,962-187,135,457 | 198495 | q35.1 - q35.2 | CN Gain | 4 | 100 | 0 | 100 |  | TLR3, FAM149A, FLJ38576, |  |
|  |  |  |  |  |  |  |  |  |  | CYP4V2 |  |
|  | chr7:142,476,116-142,486,098 | 9982 | q34 | CN Loss | 2 | 66.66666667 | 0 | 100 |  | PRSS3P2, PRSS2 | chr7:142457676, chr9:33611677 |
| Mothers | chr3:90,310,894-91,000,000 | 689106 | p11.1 - q11.1 | CN Loss | 0 | 50 | 0 | 28.14661315 |  |  | chr3:93587959 |
| Fathers | chr17:44,187,971-44,538,868 | 350897 | q21.31 | CN Gain | 7 | 37.5 | 0 | 100 | KANSL1 | KANSL1, KANSL1-AS1, LOC644172, LRRC37A, ARL17A, ARL17B, NSFP1 | chr17:45167718, chr17:43670924, chr17:45190055, chr17:66135381, chr17:30360438, chr17:62863389, chr17:44398091, chr17:44615946, chr17:43663070, chr5:140049997, chr17:65972318, chr17:65957828, chr17:62918510, chr7:25105172, chr17:65949731, chr10:27537788, chr17:37187511, chr17:30345364, chr17:29364025, chr17:66139220, chr17:30414305, chr17:37194601, chr17:28998203, chr17:43593497, chr17:45091472, chr17:62838365, chr17:66137324, chr17:30360454, chr17:43673953, chr17:44323333, chr17:28943146, chr2:148974022, chr17:28950583, chr17:66119081, chr17:30369885, chr10:27535440, chr17:29361828, chr17:28916301, chr17:43573201, chr17:41286945, chr17:41994498, chr17:44586989 |
|  | chr20:26,098,131-26,266,313 | 168182 | p11.1 | CN Gain | 2 | 37.5 | 0 | 100 |  | MIR663AHG, MIR663A | chr17:21506101, chr17:16095696, chr7:57582063, chr17:25414123, chr15:21903487, chr9:90552192, chr9:90726804, chr17:22065308, chr17:20767292, chr4:34005919, chr4:33848843, chr4:33823557, chr4:34428591, chr9:69891706, chr10:27604109, chr10:45680038, chr10:37424225, chr18:14759247, chr10:45651782, chr3:76014707, chrUn_gl000220:105861, chrUn_gl000220:149833, chr3:75976115, chr3:75920589, chr22:17225458, chr7:57714356, chr17:21895809, chr9:44817871, chr9:45049257, chr9:45784012, chr9:46170362, chr9:46330170, chr20:29572484, chr20:25850889, chr7:57873079, chr20:25781227, chr20:26031728, chr20:29419569, chr15:47398229, chr19:37524431, chr12:37987839, chrUn_gl000246:31380, chr7:57719944, chr9:67729475, chr9:44808260, chr9:45052984, chr9:45787738, chr9:46174087, chr9:46320568, chr19:37526420, chr17:21883168, chr22:17234097, chr20:26018224, chr7:57861338, chr20:29575544, chr19:40152920 |
|  | chr7:142,476,116-142,486,098 | 9982 | q34 | CN Loss | 2 | 37.5 | 0 | 100 |  | PRSS3P2, PRSS2 | chr7:142457676, chr9:33611677 |
|  | chr10:124,357,895-124,371,032 | 13137 | q26.13 | CN Loss | 1 | 37.5 | 0 | 100 | DMBT1 | DMBT1 |  |
| Hypospa | chr9:135,801,492- | 11404 | q34.13 | CN Gain | 1 | 36.36363636 | 0 | 100 | TSC1 | TSC1 |  |
| dias | 135,812,896 |  |  |  |  |  |  |  |  |  |  |
|  | chr10:43,595,893-43,618,952 | 23059 | q11.21 | CN Gain | 1 | 36.36363636 | 0 | 100 | RET | RET |  |
|  | chr3:90,310,894-91,000,000 | 689106 | p11.1 - q11.1 | CN Loss | 0 | 36.36363636 | 0 | 28.1466131 |  |  | chr3:93587959 |
|  | chr7:142,476,116-142,486,098 | 9982 | q34 | CN Loss | 2 | 54.54545455 | 0 | 100 |  | PRSS3P2, PRSS2 | chr7:142457676, chr9:33611677 |
|  | chr10:124,338,004-124,371,032 | 33028 | q26.13 | CN Loss | 1 | 36.36363636 | 0 | 100 | DMBT1 | DMBT1 |  |

On taking an aggregate of all samples with the p <0.001 we identified loss-Ch7:q34 (PRSS3P2, PRSS2) with a frequency of 38.46% (percentage of the samples in the data set having this event) and 100% of the region was covered with CNVs. On checking B/P Genes, which displays the breakpoint genes (genes that are only partially covered by the region–possible fusion sites), we identified Ch9:q34.13 (TSC1), Ch10:q11.21 (RET) and Ch10:q26.13 (DMBT1) (Table 2a).

Further using clustering algorithms all the Hypospadias samples were clustered based on similar aberration profiles. We identified five clusters; Fifth cluster (Hypospadias-5) represented all sub coronal cases (H1, H2, H19 and H27); second cluster (Hypospadias-2-H32, H37, H21), first cluster (Hypospadias-1-H9) and third cluster (Hypospadias-3-H20, H22) were related to proximal penile cases but propensity to severity increased in the order of second, first and the third cluster. Fourth cluster (Hypospadias-4) was specific to penoscrotal type (Table 3b).

**Table 3b:** Clustering of hypospadias samples based on abberation profiles. For aggregate sample p value cut off was p 0.05 and aggregate percent cutoff was 35% (listed only the regions which are lost or gained in at least 35% of the population).

| **Hypospadias-5**<br>**(H1, H2, H19,**<br>**H27)-Sub coronal** | **Region** | **Region**<br>**Length** | **Cytoband Location** | **Event** | **% of CNV Overlap** | **Gene Symbols** |
| --- | --- | --- | --- | --- | --- | --- |
|  | chr6:160,382,485-160,407,772 | 25287 | q25.3 | CN Gain | 100 | IGF2R |
|  | chr13:67,494,297-67,558,431 | 64134 | q21.32 | CN Gain | 100 | PCDH9, PCDH9-AS3 |
| **Hypospadias-2**<br>**(H32, H37, H21)-**<br>**proximal penile** | **Region** | **Region**<br>**Length** | **Cytoband Location** | **Event** | **% of CNV Overlap** | **Gene Symbols** |
|  | chr3:90,310,894-91,000,000 | 689106 | p11.1 - q11.1 | CN Loss | 28.14661 |  |
|  | chr7:142,476,116-142,486,098 | 9982 | q34 | CN Loss | 100 | PRSS3P2, PRSS2 |
| **Hypospadias-1**<br>**(H9)-Proximal**<br>**penile** | **Region** | **Length** | **Cytoband** | **Event** | **% of CNV Overlap** | **Gene Symbols** |
|  | chr3:90,310,894-91,000,000 | 689107 | p11.1 - q11.1 | CN Loss | 28.14661 |  |
|  | chr6:61,000,000-62,241,352 | 1241353 | q11.1 | CN Loss | 29.09336 |  |
|  | chr7:142,476,116-142,486,098 | 9983 | q34 | Homozygous<br>Copy Loss | 100 | PRSS3P2, PRSS2 |
|  | chr8:2,179,612-2,493,957 | 314346 | p23.3 - p23.2 | CN Gain | 100 | LOC101927815 |
|  | chr9:135,800,034-135,814,561 | 14528 | q34.13 | CN Gain | 100 | TSC1 |
|  | chr10:43,595,893-43,623,186 | 27294 | q11.21 | CN Gain | 90.34551 | RET |
|  | chr11:34,250,943-34,623,732 | 372790 | p13 | CN Gain | 85.6844 | ABTB2, CAT, ELF5 |
|  | chr11:54,827,208-55,030,580 | 203373 | q11 | CN Loss | 100 | TRIM48 |
|  | chr17:44,045,984-44,050,801 | 4818 | q21.31 | CN Loss | 100 | MAPT |
|  | chr22:23,577,112-23,579,423 | 2312 | q11.23 | Homozygous<br>Copy- Loss | 100 | BCR |
| **Hypospadias-3**<br>**(H20, H22)-**<br>**Proximal Penile** | **Region** | **Region**<br>**Length** | **Cytoband Location** | **Event** | **% of CNV Overlap** | **Gene Symbols** |
|  | chr1:35,637,150-35,677,978 | 40828 | p34.3 | CN Gain | 61.69785 | SFPQ |
|  | chr1:248,651,682-248,794,849 | 143167 | q44 | CN Loss | 100 | OR2T5, OR2G6, OR2T29,<br>OR2T34, OR2T10, OR2T11 |
|  | chr3:30,660,335-30,697,591 | 37256 | p24.1 | CN Gain | 100 | TGFBR2 |
|  | chr3:70,006,991-70,054,976 | 47985 | p13 | CN Gain | 4.436803 | MITF, LINC01212 |
|  | chr3:71,106,700-71,181,285 | 74585 | p13 | CN Gain | 100 | FOXP1 |
|  | chr3:90,109,552-90,182,046 | 72494 | p11.1 | CN Gain | 100 |  |
|  | chr4:87,868,929-87,966,793 | 97864 | q21.3 | CN Gain | 22.5844 | AFF1 |
|  | chr4:181,855,644-181,983,685 | 128041 | q34.3 | CN Gain | 100 |  |
|  | chr5:4,908,739-5,022,241 | 113502 | p15.32 | CN Gain | 50.82994 |  |
|  | chr5:50,016,088-50,048,299 | 32211 | q11.1 | CN Gain | 100 | PARP8 |
|  | chr5:67,505,065-67,596,451 | 91386 | q13.1 | CN Gain | 68.62867 | PIK3R1 |
|  | chr6:26,009,559-26,474,214 | 464655 | p22.2 | CN Gain | 97.23042 | HIST1H1A, HIST1H3A,<br>HIST1H4A, HIST1H4B,<br>HIST1H3B, HIST1H2AB,<br>HIST1H2BB, HIST1H3C,<br>HIST1H1C, HFE, HIST1H4C,<br>HIST1H1T, HIST1H2BC,<br>HIST1H2AC, HIST1H1E,<br>HIST1H2BD, HIST1H2BE,<br>HIST1H4D, HIST1H3D,<br>HIST1H2AD, HIST1H2BF,<br>HIST1H4E, HIST1H2BG,<br>HIST1H2AE, HIST1H3E,<br>HIST1H1D, HIST1H4F,<br>HIST1H4G, HIST1H3F,<br>HIST1H2BH, HIST1H3G,<br>HIST1H2BI, HIST1H4H,<br>BTN3A2, BTN2A2, BTN3A1,<br>BTN2A3P, BTN3A3,<br>BTN2A1, LOC285819 |
|  | chr6:27,727,871-27,868,980 | 141109 | p22.1 | CN Gain | 86.70956 | LOC100131289, HIST1H2BL,<br>HIST1H2AI, HIST1H3H,<br>HIST1H2AJ, HIST1H2BM,<br>HIST1H4J, HIST1H4K,<br>HIST1H2AK, HIST1H2BN,<br>HIST1H2AL, HIST1H1B,<br>HIST1H3I, HIST1H4L,<br>HIST1H3J, HIST1H2AM,<br>HIST1H2BO |
|  | chr6:160,378,263-160,411,195 | 32932 | q25.3 | CN Gain | 100 | IGF2R |
|  | chr7:62,437,464-62,510,943 | 73479 | q11.21 | CN Gain | 100 |  |
|  | chr7:77,240,174-77,549,202 | 309028 | q11.23 - q21.11 | CN Gain | 40.17791 | PTPN12, APTR, RSBN1L, TMEM60, PHTF2 |
|  | chr7:142,476,116-142,486,098 | 9982 | q34 | CN Loss | 100 | PRSS3P2, PRSS2 |
|  | chr9:135,800,034-135,812,896 | 12862 | q34.13 | CN Gain | 100 | TSC1 |
|  | chr10:43,590,678-43,618,952 | 28274 | q11.21 | CN Gain | 100 | RET |
|  | chr10:89,611,321-89,698,354 | 87033 | q23.31 | CN Gain | 28.0836 | KLLN, PTEN |
|  | chr10:124,338,004-124,382,102 | 44098 | q26.13 | CN Loss | 100 | DMBT1 |
|  | chr11:5,785,663-5,808,947 | 23284 | p15.4 | CN Loss | 100 | OR52N5 |
|  | chr11:81,502,443-81,519,577 | 17134 | q14.1 | CN Loss | 100 |  |
|  | chr11:128,575,283-128,613,047 | 37764 | q24.3 | CN Gain | 100 | FLI1 |
|  | chr12:65,577,673-65,609,172 | 31499 | q14.3 | CN Gain | 100 | LEMD3 |
|  | chr13:67,494,297-67,562,997 | 68700 | q21.32 | CN Gain | 100 | PCDH9, PCDH9-AS3 |
|  | chr17:21,157,940-21,198,074 | 40134 | p11.2 | CN Gain | 24.08183 | MAP2K3 |
|  | chr18:48,562,070-48,579,356 | 17286 | q21.2 | CN Gain | 0 | SMAD4 |
|  | chr19:54,638,178-54,714,174 | 75996 | q13.42 | CN Gain | 100 | CNOT3, LENG1, TMC4, MBOAT7, TSEN34, RPS9 |
|  | chr20:29,932,833-29,982,244 | 49411 | q11.21 | CN Gain | 100 | DEFB118, DEFB119 |
|  | chr20:62,764,372-62,768,514 | 4142 | q13.33 | CN Loss | 100 |  |
|  | chr21:38,756,746-38,808,477 | 51731 | q22.13 | CN Gain | 100 | DYRK1A |
| **Hypospadias-4 (H7, H38)-Penoscrotal** | **Region** | **Region Length** | **Cytoband Location** | **Event** | **% of CNV Overlap** | **Gene Symbols** |
|  | chr3:90,310,894-91,000,000 | 689106 | p11.1 - q11.1 | CN Loss | 28.14661 |  |
|  | chr4:168,481,367-168,651,682 | 170315 | q32.3 | CN Loss | 100 |  |
|  | chr7:142,476,116-142,486,098 | 9982 | q34 | CN Loss | 100 | PRSS3P2, PRSS2 |
|  | chr8:39,238,717-39,386,527 | 147810 | p11.22 | CN Gain | 100 | ADAM5, ADAM3A |
|  | chr8:78,615,277-78,808,102 | 192825 | q21.12 | CN Loss | 75.98392 |  |
|  | chr9:135,800,034-135,812,896 | 12862 | q34.13 | CN Gain | 100 | TSC1 |
|  | chr10:962,366-1,210,801 | 248435 | p15.3 | CN Gain | 100 | LARP4B, LOC101927762, GTPBP4, IDI2, IDI2-AS1, IDI1, WDR37, LINC00200 |
|  | chr10:43,595,893-43,645,508 | 49615 | q11.21 | CN Gain | 74.50368 | RET, CSGALNACT2 |
|  | chr10:124,338,004-124,371,032 | 33028 | q26.13 | CN Loss | 100 | DMBT1 |
|  | chr11:471,611-516,933 | 45322 | p15.5 | CN Loss | 100 | PTDSS2, RNH1 |
|  | chr16:34,506,306-34,724,224 | 217918 | p11.2 - p11.1 | CN Gain | 100 | LOC283914, LOC146481 |
|  | chr17:34,431,863-34,649,973 | 218110 | q12 | CN Gain | 100 | CCL4, CCL3L3, CCL3L1, CCL4L1, CCL4L2, TBC1D3H, LOC101060321, TBC1D3F, CCL3L3, CCL3L1, CCL4L1, CCL4L2 |
|  | chr22:25,691,453-25,909,031 | 217578 | q11.23 - q12.1 | CN Gain | 100 | IGLL3P, LRP5L, CRYBB2P1, MIR6817 |

On analyzing the samples through ChAS software we identified loss of Ch: 7-PRSS3P2, PRSS2 (CN=0.00) and Ch: 16-CES1P2, CES1P1 (CN=1.00) in 50% and 41.6% cases respectively. Gain of Ch: X-AVPR2, ARHGAP4, NAA10 (CN=2.00) and XG (CN=2.00) genes was observed in about 58.33% and 41.6% respectively. Ch: Y mosaicism (1.05-1.11) was reported in 75% cases. However, 33.33% cases reported the loss of TMPRSS11E, UGT2B17, UGT2B15 (CN=1.00) genes in Ch: 4 and gain of PRKCZ, C1orf86, SKI, MORN1 (CN=3.00), PRDM16, ARHGEF16, MEGF6, MIR551A, TPRG1L, WRAP73, TP73 (CN=3.00) genes in Ch: 1. Rest of the variants were detected in 16.66% of hypospadias sample (Table 4). Also when compared with UCSC public decipher database, which catalogs genetic abnormalities i.e. CNVs with phenotypic data, we identified some variants which corroborated with the previously reported hypospadias and its associated anomalies as shown in (Table 4).

**Table 4:**
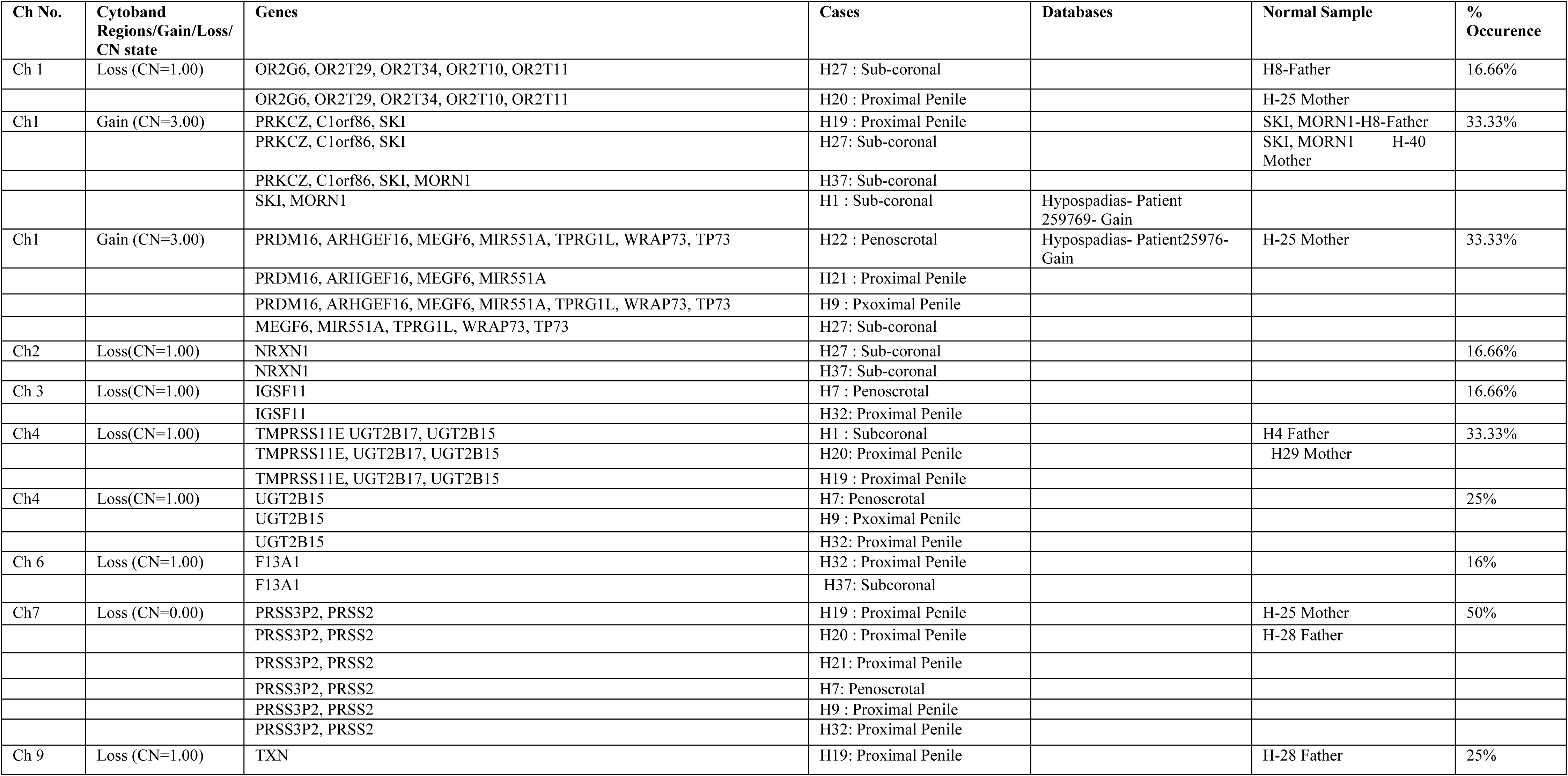
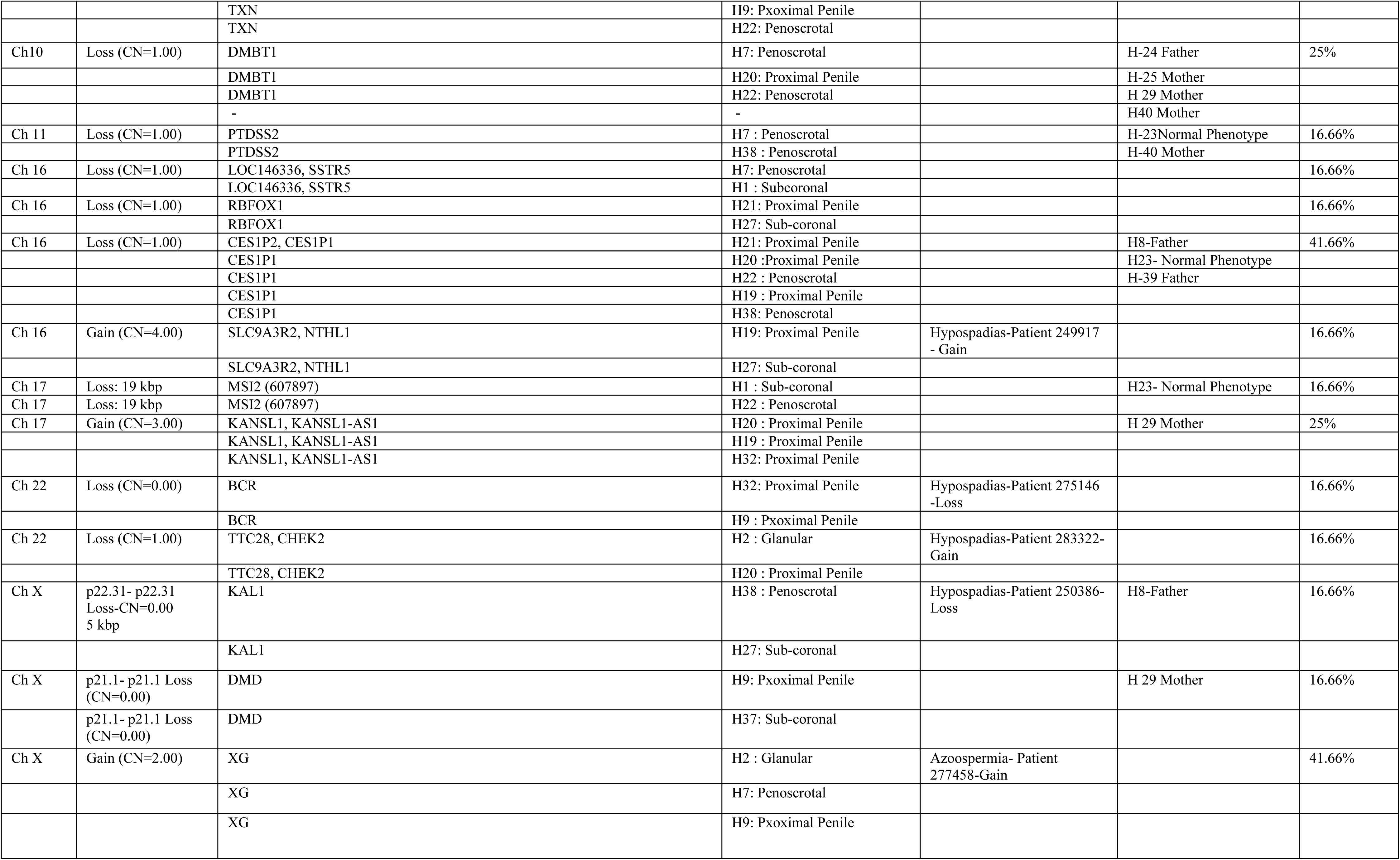
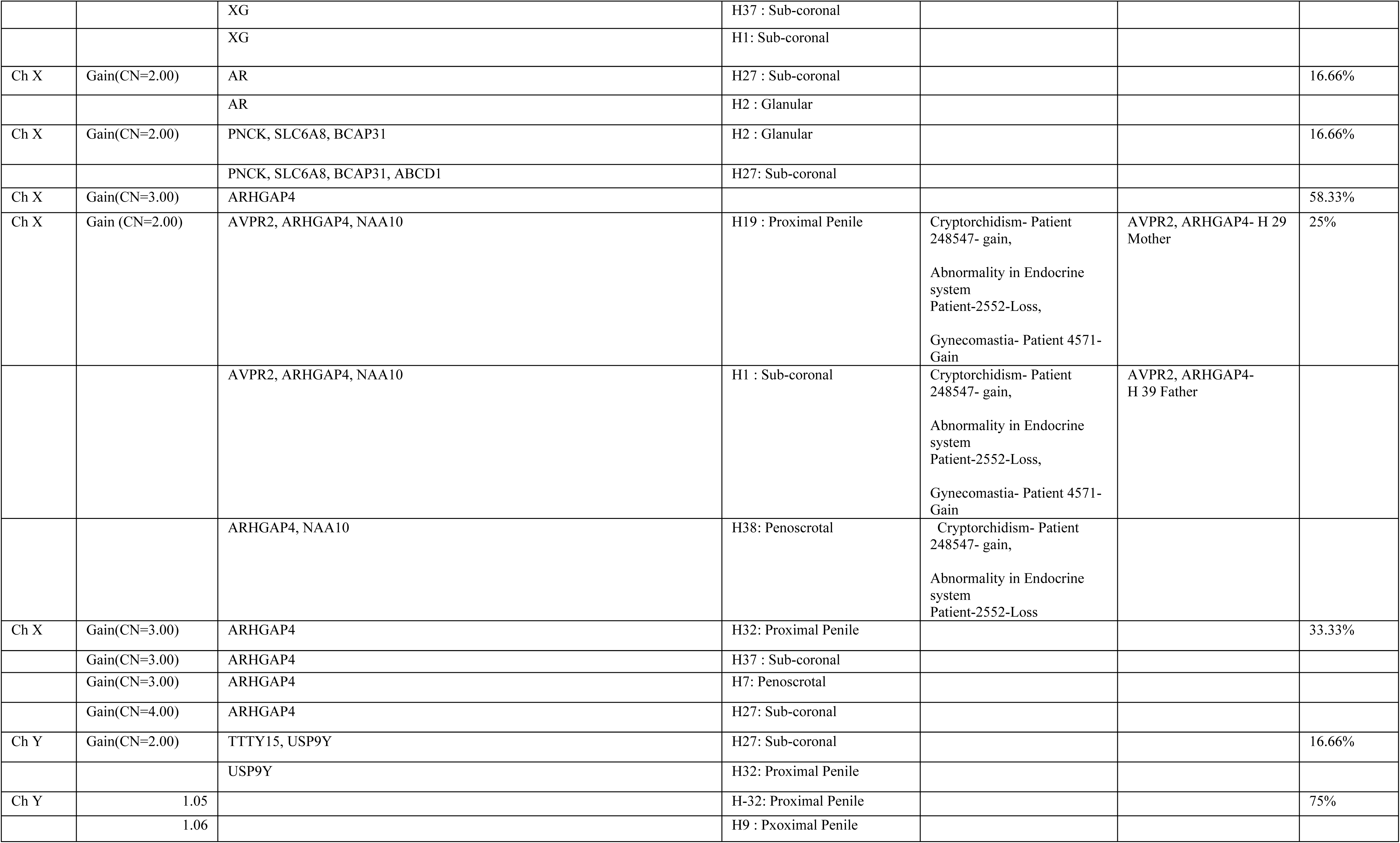
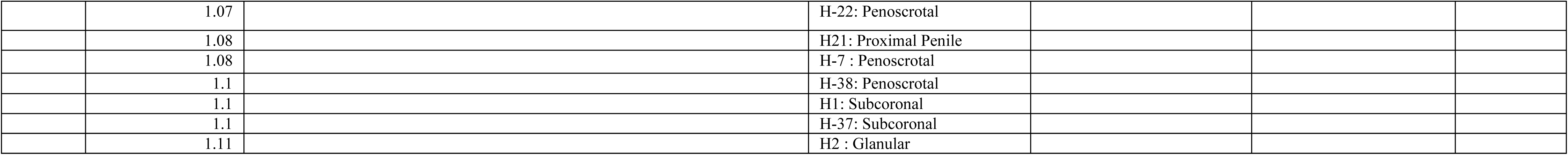
Percent occurrence of Copy Number Variations including multiple genes in all Hypospadias as well as presence of same regions in Normal Samples identified through ChAS.

### Validation of the copy number variation by TaqMan copy number assay

Using the probe against target sequence: PRSS3P2 and reference sequence TERT a total of 33 samples were processed for identification of copy number assay through Taqman based assay. Out of 33 samples, one was negative control and remaining 32 were hypospadias samples. Out of 32, 13 samples showed 0 Copy, three samples showed 1 Copy, 13 samples showed 2 Copies and one sample showed −3 Copies with the ΔCт variability of 0.1666 (Table 5).

**Table 5:** The copy numbers of the PRSS3P2 genes in 32 hypospadias samples were detected through TaqMan® Copy Number Assays and measured by CopyCaller™ Software v1.0.

| **Sample ID** | **Type of hypospadias** | **copy number calculated** | **copy number predicted** | **confidence** |
| --- | --- | --- | --- | --- |
| **AH01** | **Distal penile hypospadias** | **1.48** | **2.0** | **0.97** |
| **AH02** | **Proximal penile hypospadias** | **0.73** | **0** | **-** |
| **AH03** | **Distal penile hypospadias** | **0.9** | **0** | **-** |
| **AH04** | **Distal penile hypospadias** | **1.53** | **2** | **0.99** |
| **AH05** | **Distal penile hypospadias** | **0.43** | **0** | **-** |
| **AH06** | **Distal penile hypospadias** | **0.74** | **0** | **-** |
| **AH07** | **Hypospadias** | **1.24** | **1** | **0.455** |
| **AH08** | **Hypospadias** | **0.69** | **0** | **-** |
| **AH09** | **Hypospadias** | **0.69** | **0** | **-** |
| **AH10** | **Distal penile hypospadias** | **0.71** | **0** | **-** |
| **AH11** | **Coronal Hypospadias** | **2.58** | **3** | **0.455** |
| **AH12** | **Coronal Hypospadias** | **1.93** | **2** | **0.99** |
| **H1** | **Sub-coronal Hypospadias** | **2.07** | **2** | **0.97** |
| **H2** | **Glanular** | **1.68** | **2** | **0.995** |
| **H7** | **Penoscrotal with chordee** | **0.82** | **0** | **-** |
| **H9** | **Proximal penile hypospadias** | **1.05** | **0** | **-** |
| **H11** | **Proximal penile hypospadias** | **1.07** | **0** | **-** |
| **H13** | **Glanular Hypospadias** | **2.11** | **2** | **0.97** |
| **H15** | **Proximal penile hypospadias** | **0.96** | **0** | **-** |
| **H16** | **Coronal Hypospadias** | **2.04** | **2** | **0.99** |
| **H17** | **Proximal penile** **hypospadias** | **0.97** | **0** | **-** |
| **H18** | **Sub-coronal hypospadias** | **1.96** | **2** | **0.99** |
| **H19** | **Proximal penile hypospadias** | **1.88** | **2** | **0.95** |
| **H20** | **Proximal penile hypospadias** | **0.18** | **0** | **-** |
| **H21** | **Proximal penile hypospadias** | **1.95** | **2** | **0.99** |
| **H22** | **Proximal penile hypospadias** | **2.15** | **2** | **0.95** |
| **H27** | **Sub-coronal hypospadias** | **2.1** | **2** | **0.96** |
| **H32** | **Proximal penile hypospadias** | **1.24** | **1** | **0.455** |
| **H33** | **Sub-coronal hypospadias** | **2.11** | **2** | **0.95** |
| **H34** | **Glanular hypospadias** | **1.78** | **2** | **0.995** |
| **H37** | **Proximal penile hypospadias** | **1.22** | **1** | **0.62** |
| **H38** | **Penoscrotal hypospadias** | **1.05** | **0** | **-** |

## Discussion

Until date, several studies were conducted on Hypospadias, for a genome-wide association study. Geller *et al.,* (Geller et al. 2014), has been tested on hypospadias cases and control, and identified 18 genomic regions showing independent association with *P* < 5 × 10^−8^ with probability of 9% to develop this condition. (Carmichael et al. 2014) Carmichael *et al.,* suggested the role of foetal sex hormone biosynthesis and metabolism pathway in hypospadias for association between lower testosterone or DHT levels or higher oestrogen or progesterone levels. Various isolated Hypospadias studies have been conducted to show association with the Hypospadias-risk. Most importantly, Diacylglycerol Kinase Kappa (*DGKK*6) which showed differential genetic association with hypospadias risk, positive in Caucasians and negative in the Chinese population. Indian reports have also shown significant association between SRD5A2-V89L (Samtani et al. 2011), SRD5A2-A49T, R227Q and TA repeat gene polymorphisms (Samtani et al. 2015) and MAMLD1 (c.2960C>T) (Ratan et al. 2016) with hypospadias risk and also the severity of the disease. Hence, till date, few genetic associations have been reported in Indian studies but no comprehensive approach has been followed to assess the genetic involvement of different SNPs contributing to pathways like sex hormone biosynthesis and metabolism; embryonic development and Phospholipase D signaling with the severity of Hypospadias. Although besides SNPs, CNVs that involve genomic segments containing one or more genes can result in several genetic disorders and complex diseases.

Hence, to establish a correlation between the severity of Hypospadias and significant SNPs/gain of a particular region/loss of a particular region we performed this study. In our report, all the genetic variants related to embryonic development pathway were not related to the severity of the disease. Although, sex hormone biosynthesis and metabolism gene-variants were involved with the severity of the disease. Association of rs17268974 of Steroid Sulfatase-STS gene (Microsomal) suggests an alteration in catalyzing ability of the gene product in the conversion of sulfated steroid precursor to the generation of placental estrogens.

In Penoscrotal type of hypospadias, the involvement of all the seven SNPs (rs5934740; rs5934842; rs5934913; rs6639811; rs3923341; rs17268974; rs5934937) of STS gene further confirms the hampering of biosynthesis of oestrogens or oestriols. However, all the seven STS-SNPs have been associated with risk and not to the severity (Carmichael et al. 2014).

Further, besides oestrogens alteration of conversion of testosterone to dihydrotestosterone is also suggestive through association of rs7562326 of SRD5A2-Steroid 5-alpha-reductase in the urethral seam. Although SRD5A2-genetic variants have been associated with hypospadias in several small studies (Makridakis et al. 2000; Thai et al. 2005; Sata et al. 2010; Samtani et al. 2011) but not in one larger study (van der Zanden et al. 2010).

Association of rs1877031 of STARD3 gene-STAR-related lipid transfer-also supports the development of Penoscrotal Hypospadias by conversion of sulfated steroid precursors to Oestrogen. Both STARD3 SNPs (rs1874224; rs1877031) have been shown to be associated with risk; results varying between whites and Hispanics and by severity also (Carmichael et al. 2014). Hence, compared to Glanular+Coronal, Penile/Penoscrotal are associated with increased number of sex hormone biosynthesis and metabolism gene-variants SNPs i.e. severity of disease

Significantly, we observed loss of Ch3:p11.1 - q11.1 region of maternal origin and loss of Ch7:q34 (PRSS3P2, PRSS2) as well as Ch10:q26.13-loss (DMBT1) of paternal origin in Hypospadias cases. However, gain of Ch9:q34.13 (TSC1) and Ch10:q11.21 (RET) was *de*-*novo* to Hypospadias samples. The identified variant Ch7:q34 (PRSS3P2, PRSS2) was flanked with segmental duplication of chr9:33611677 which may allow recombination by non-allelic homologous recombination as suggested (Coughlin et al. 2012; Shaikh et al. 2000; Shaikh et al. 2007). However, through breakpoint analysis we identified possible fusion sites in Ch9:q34.13-TSC1, Ch10:q11.21-RET and Ch10:q26.13--DMBT1 where non-homologous end joining and replication-based mechanisms such as fork stalling and template switching can occur (van Binsbergen 2011; Korbel et al. 2007).

Functionally, the complete loss of protease, serine, 3 pseudogene 2PRSS3P2 (five exons) in hypospadias samples, which encode a protein similar to trypsinogen, localized to the T cell receptor beta locus on chromosome 7 may alter probably the cell migration (PRSS3P2 gene card). Further, PRSS3P2 showed concordance with gain of TSC1 (tuberous sclerosis 1) a protein-coding gene encoding a growth inhibitory protein playing role in the stabilization of tuberin. Subsequently TSC1 showed concordance with gain of RET, a member of the cadherin superfamily, encoding one of the receptor tyrosine kinases, and are cell-surface molecules that transduce signals for cell growth and differentiation and loss of DMBT1 (deleted in malignant brain tumors 1), a protein-coding gene involved in mucosal defense system, cellular immune defense and epithelial differentiation (Fig.1a and b). Importantly, the gains and losses observed in our hypospadias samples have not been reported previously in decipher database. Through cluster analysis we observed that PRSS3P2 was not observed in fifth-cluster linked to subcoronal cases, however, the predisposition to hypospadias increased with the homozygous loss of PRSS3P2. Finally, in penoscrotal cases besides all the other variants loss of DMBT1 was also observed (Table 6). The observations identified are quite promising; however, on further validation in previously copy number characterized 12-hypospadias cases, we identified that subcoronal samples (Hypospadias cluster-5) did not show any copy number deletion of PRSS3P2 through Taqman assay. While, rest of the clusters (Hypospadias 2, 1, 3 and 4) depicted 1 or 2 copies loss except H22 which showed the presence of both the copies of PRSS3P2 gene (Table 5). Additionally, in 20-new samples, most of the grade 3 (distal penile, proximal penile and penoscrotal) hypospadias showed loss of PRSS3P2 while grade1 and grade 2 (glanular coronal and subcoronal) hypospadias represented no copy number loss (Table 5).

**Fig. 1a.**
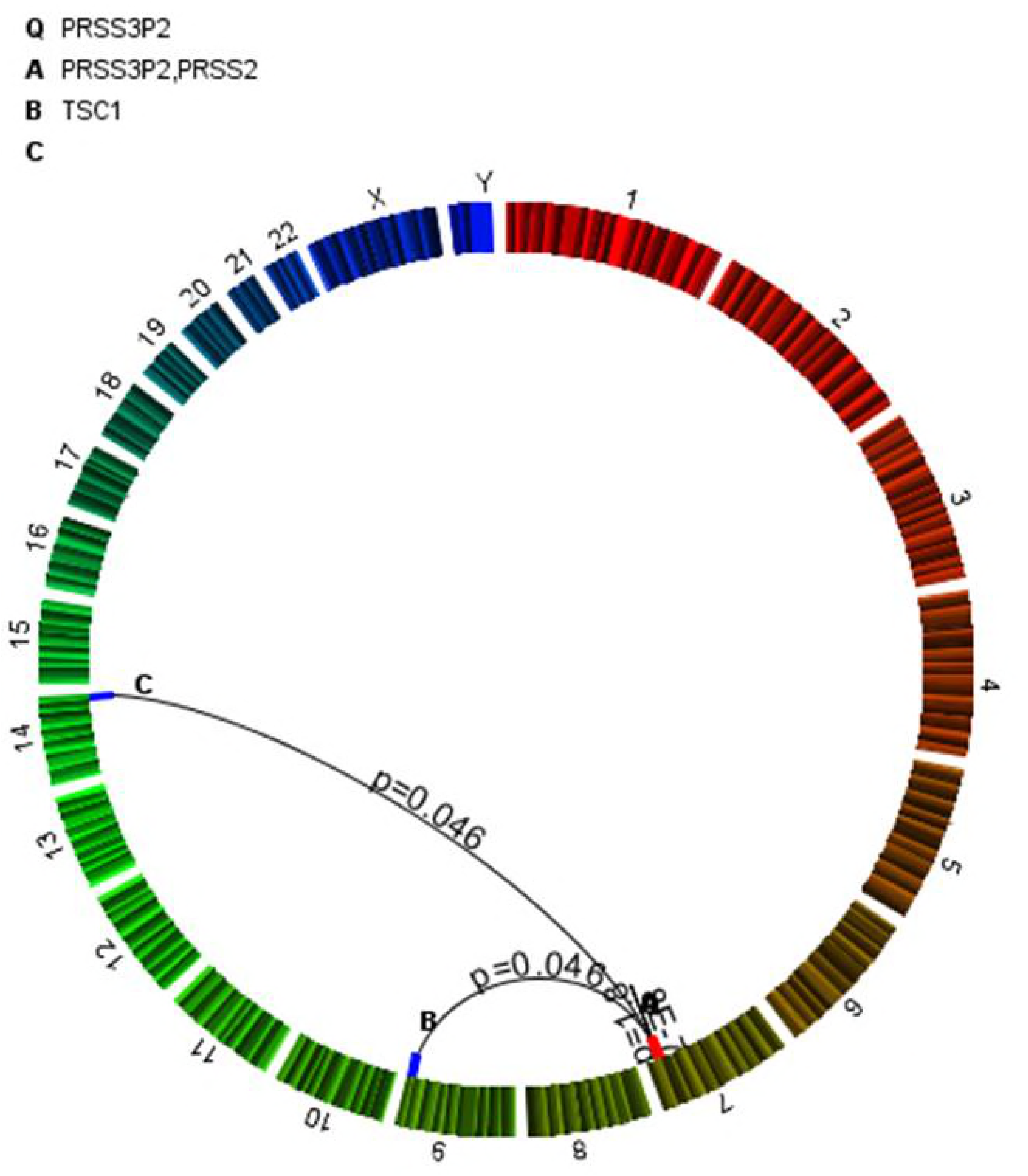
Concordance of PRSS3P2

**Fig. 1b.**
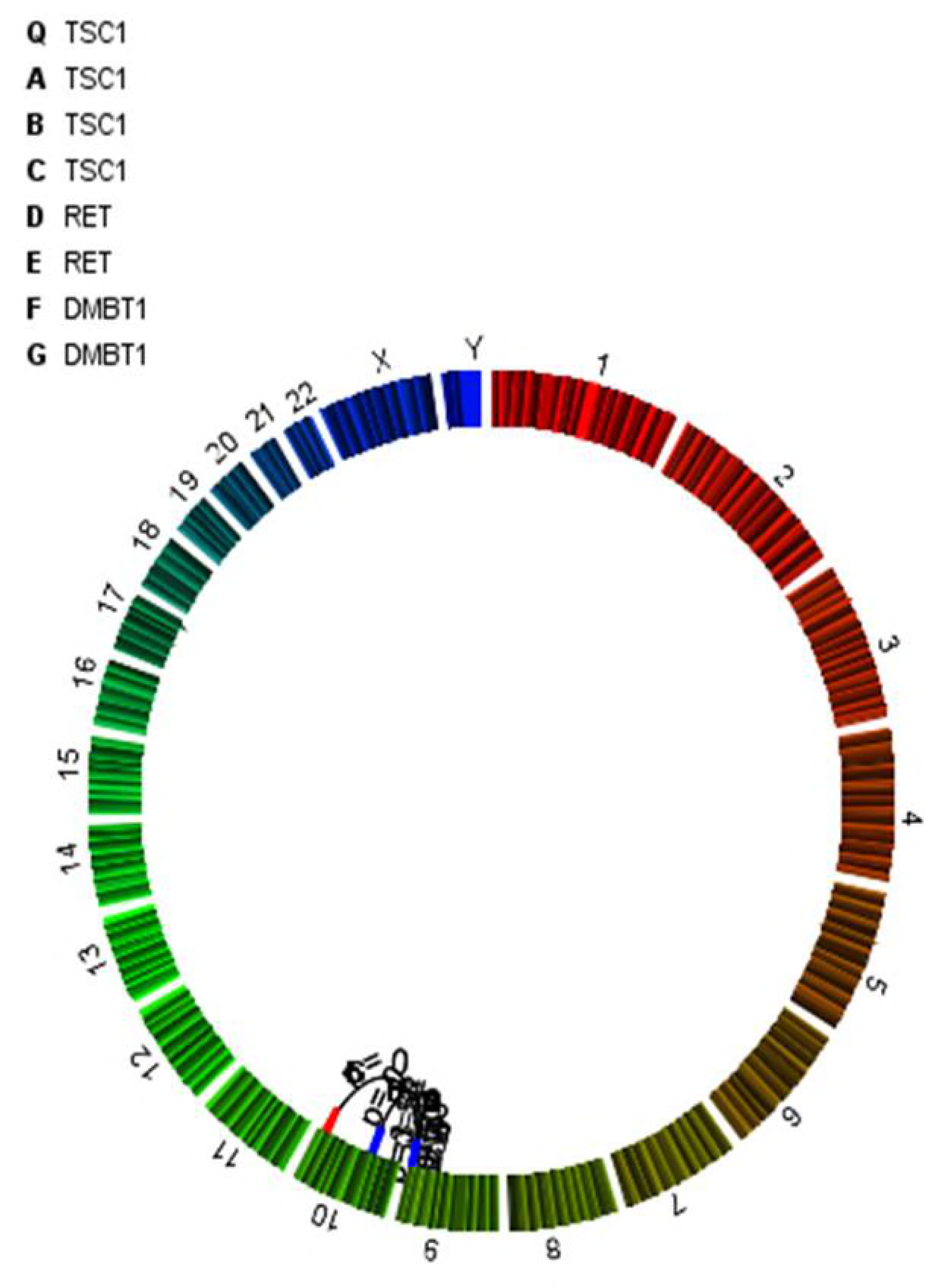
Concordance of TSCl

**Table 6:**
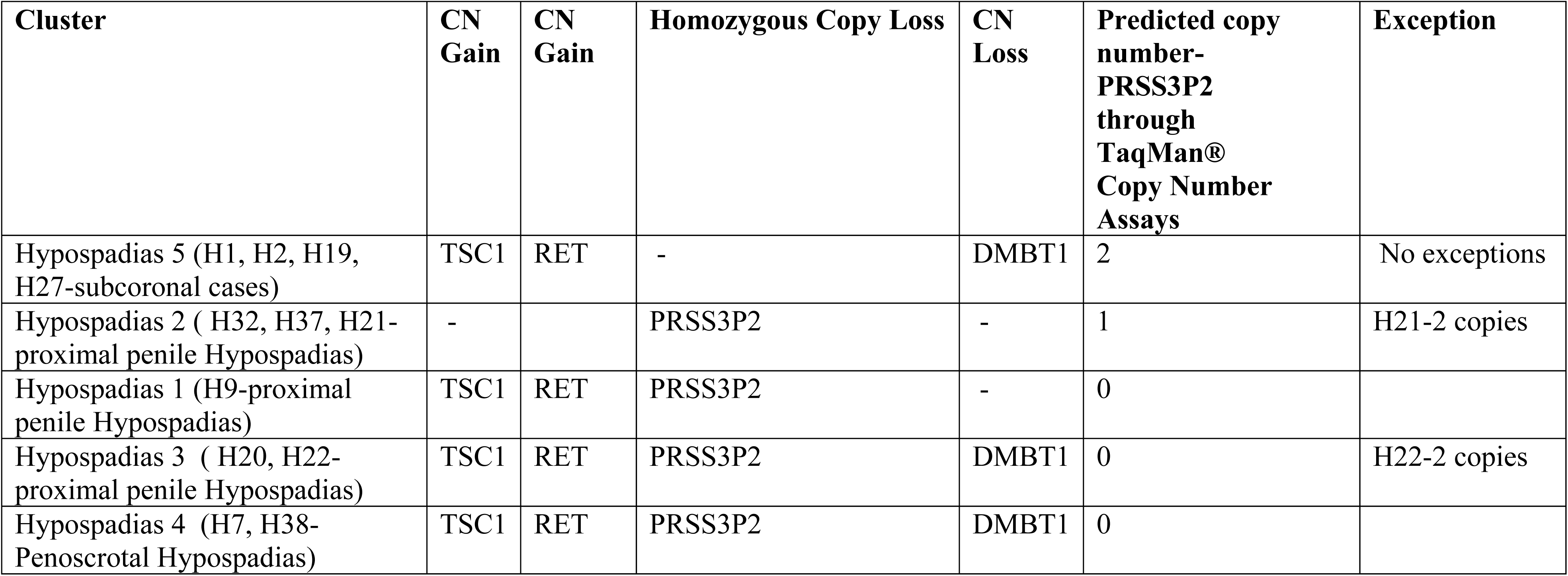
Significantly identified genes in various clusters. Additionally, copy number deletion of PRSS3P2 gene identified through aCGH method was validated by TaqMan® Copy Number Assays.

| **Cluster** | **CN Gain** | **CN Gain** | **Homozygous Copy Loss** | **CN Loss** | **Predicted copy number-PRSS3P2 through TaqMan® Copy Number Assays** | **Exception** |
| --- | --- | --- | --- | --- | --- | --- |
| Hypospadias 5 (H1, H2, H19, H27-subcoronal cases) | TSC1 | RET | - | DMBT1 | 2 | No exceptions |
| Hypospadias 2 ( H32, H37, H21-proximal penile Hypospadias) | - |  | PRSS3P2 | - | 1 | H21-2 copies |
| Hypospadias 1 (H9-proximal penile Hypospadias) | TSC1 | RET | PRSS3P2 | - | 0 |  |
| Hypospadias 3 ( H20, H22-proximal penile Hypospadias) | TSC1 | RET | PRSS3P2 | DMBT1 | 0 | H22-2 copies |
| Hypospadias 4 (H7, H38-Penoscrotal Hypospadias) | TSC1 | RET | PRSS3P2 | DMBT1 | 0 |  |

## Conclusion

In this study, we identified and reported novel copy number variation (CNVs; gains and losses) & mosaicism in the twenty-four cases of hypospadias (both confirmed Hypospadias cases and control (parental) samples) which have not been reported previously in decipher database. we have identified loss of Ch7:q34 (PRSS3P2, PRSS2) as a novel locus for hypospadias which also shows concordance with TSC1-gain, RET-gain, and DMBT1-loss. Hence, we can suggest that Grade 1 and 2 (including sub coronal and glanular) hypospadias cases show no loss of PRSS3P2 with no association of STS; SRD5A2; STARD3-gene but in Grade 3 and 4 (including Penile and Penoscrotal) hypospadias cases loss of PRSS3P2 gene accompanied by association of STS; SRD5A2; STARD3 was observed. Although gain of TSC1, RET, and loss of DMBT-1 was observed in both the Grades. Hence, we conclude that the homozygous loss of PRSS3P2 accompanied with the association of STS; SRD5A2; STARD3 may link to the severity of the disease.

## Acknowledgements

This work was supported by University Grant from King George Medical University, Lucknow, India. No funding sources are involved for accomplishment of this study. We are grateful to Dr. Paras Yadav, Senior Application Scientist Imperial Life Sciences for technical assistance. Previously, the abstract has been submitted to “European Paediatric Surgeons’ Association-15th Congress Dublin, Ireland-2014 on 18-21st June 2014”.

